# Multiple mechanisms of photoreceptor spectral tuning following loss of UV color vision in *Heliconius* butterflies

**DOI:** 10.1101/2021.02.23.432523

**Authors:** Kyle J. McCulloch, Aide Macias-Muñoz, Ali Mortazavi, Adriana D. Briscoe

## Abstract

Color vision modifications occur in animals via a process known as spectral tuning. In *Heliconius* butterflies, a genus-specific *UVRh* opsin duplication led to the evolution of UV color discrimination in *Heliconius erato* females, a rare trait among butterflies. In the *H. melpomene* and *H. ismenius* lineages, the UV2 receptor has been lost. Here we compare how loss of the UV2 photoreceptor has altered the visual system of these butterflies. We compare visual system evolution in three *Heliconius* butterfly species using a combination of intracellular recordings, ATAC-seq, and antibody staining. We identify several spectral tuning mechanisms including adaptive evolution of opsins, deployment of two types of filtering pigments, and co-expression of two distinct opsins in the same cell. Our data show that opsin gain and loss is driving rapid divergence in *Heliconius* visual systems via tuning of multiple spectral classes of photoreceptor in distinct lineages, potentially contributing to ongoing speciation in this genus.

## Introduction

The visual systems of arthropods and vertebrates have evolved to discriminate many distinct colors, complementing the salient visual signals found in the animal’s visual space (Briscoe and Chittka, 2001; Futahashi et al., 2015; Osorio and Vorobyev, 2005). In animals that inhabit and sense colorful environments, the evolution of new spectral channels—mainly achieved through duplication of opsin genes followed by sequence divergence to generate novel photoreceptor cell types—has been the focus of most research (Briscoe and Chittka, 2001; Chen et al., 2016; Cronin et al., 1996; Frentiu et al., 2007; Osorio and Vorobyev, 2008; Parry et al., 2005; Thoen et al., 2014). Opsin proteins covalently bond a vitamin A-derived chromophore to make rhodopsin, the visual pigment responsible for detection of light in animal photoreceptor cells (Palczewski et al., 2000). Among opsin duplicates, amino acid substitutions that interact with the chromophore may shift the probability that a particular rhodopsin absorbs a photon of light at a given wavelength (Arshavsky et al., 2002; Bloch, 2016; Shichida and Matsuyama, 2009). Other ways of modifying a photoreceptor neuron’s sensitivity to light include the addition of photostable filtering pigments and co-expression of multiple opsins in a single cell (Arikawa et al., 2003; K. Arikawa et al., 1999; Kentaro Arikawa et al., 1999; Dalton et al., 2014; Knott et al., 2010; Satoh et al., 2017; Vöcking et al., 2017; Wakakuwa et al., 2004).

Despite the apparent ease with which opsins duplicate, most animals have four or fewer spectral channels, perhaps because of the increasing complexity of the neural wiring required to add spectrally opponent channels to the nervous system (Barlow, 1981; Buchsbaum and Gottschalk, 1983; Dacey, 1999; Kelber and Osorio, 2010; Osorio, 1986; Rushton, 1972; Thornton and Pugh, 1983). Given constraints on adding more spectral types of photoreceptor, such additions may not always be adaptive. Shifting selection pressures or drift might result in the loss of spectral channels, which could be beneficial, even in a colorful world. Loss of visual capability is generally studied in the context of low light environments where color vision is no longer necessary, such as in shifts to nocturnality, caves, or deep ocean. We decided to address how photoreceptor loss might affect the molecular and physiological evolution of *Heliconius* visual systems, butterflies with no apparent loss in their color sensing demands.

Butterflies in the genus *Heliconius* have superb color vision from short UV to near-infrared wavelengths (Finkbeiner et al., 2017; Finkbeiner and Briscoe, 2020; McCulloch et al., 2016a; Zaccardi et al., 2006). The morphological basis of *Heliconius* color vision is the compound eye, which is similar in structure to other butterflies. The eye is a retinal mosaic of thousands of unit eyes, called ommatidia. Each ommatidium is a long tube of nine photoreceptor cells that project axons to the optic lobe (Figure 1A). The photoreceptor cells (R1-9) project microvilli packed with rhodopsin into the center of the ommatidium, forming a fused fiber optic-like structure called the rhabdom. In transverse sections, the R1-8 cells in a single ommatidium are arranged like petals on a flower with the rhabdom in the center (Figure 1A). Light is focused through the cornea and crystalline cone and channeled through the rhabdom where it is absorbed by rhodopsins of each photoreceptor cell type. R1 and R2 cells are variable short-wavelength photoreceptor cells (BRh- or UVRh-expressing), involved in color vision (Borst et al., 2010; Schnaitmann et al., 2018). The R3-8 cells express LWRh, are green-sensitive, and are involved in motion and contrast vision, though a subset may also contribute to color processing (Chen et al., 2020). A subset of R5-8 cells have an orange pigment next to the proximal rhabdom, a filtering pigment responsible for yellow-to-red color vision in *Heliconius* (Figure 1A) (Zaccardi et al., 2006).

**Fig. 1.**
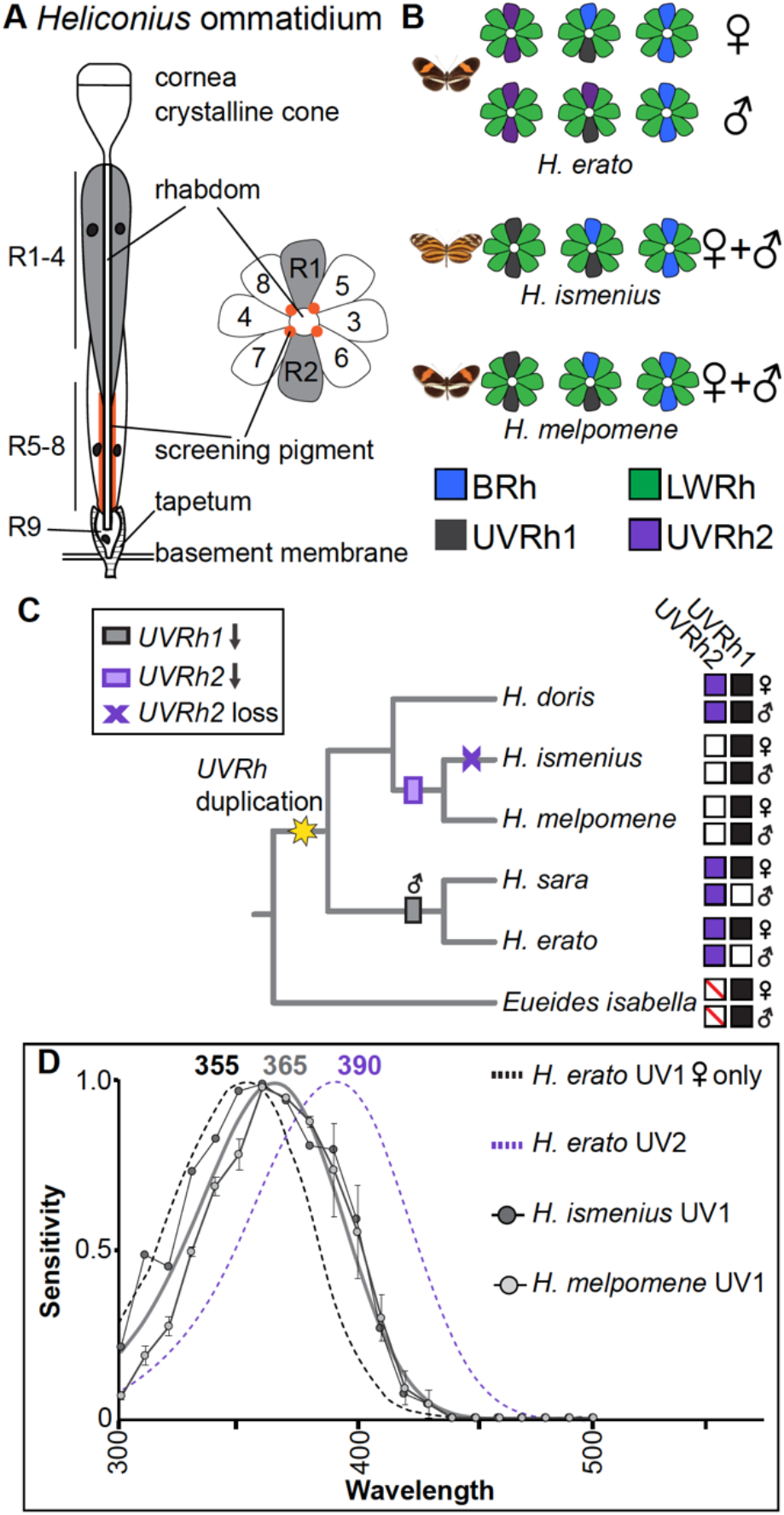
*Heliconius* eye anatomy and UVRh evolutionary trajectory and physiology. **A)** Overview of *Heliconius* ommatidium structure. Light enters and is focused through the cornea and crystalline lens and passes through the rhabdom. R1-R4 cells contribute more distally to the rhabdom while R5-8 cells and filtering pigment (when present) contribute more proximally. The tiny R9 cell sits at the proximal end of the rhabdom. The cells are supplied with oxygen via the tracheal tapetum that penetrates through the basement membrane and surrounds the proximal end of the ommatidium. Cell axons project through the basement membrane to the optic lobe. In transverse sections, R1/R2 cells sit opposite each other while R5-8 have orange pigment next to the rhabdom from around 320-480 μm below the cornea (Zaccardi et al. 2006). The density of this pigment is heterogeneous between ommatidia across the eye, producing yellow- and red-reflecting adjacent ommatidia. **B)** Cross sections of ommatidial types found in the compound eyes of *H. erato*, *H. melpomene*, and *H. ismenius*, based on previous antibody staining (see new results with expanded ommatidial types in Fig. 5 and Fig. S2) (McCulloch et al., 2017). Ommatidial types are classified by the combination of short wavelength opsins expressed in R1/R2. **C)** Phylogeny of major *Heliconius* groups with *Eueides* outgroup lacking the *UVRh* duplication. The *Heliconius UVRh* duplication occurred at the base of the genus, which gave rise to functional UVRh1 and UVRh2 opsins in the *Heliconius* radiation. Subsequent loss of mRNA expression is mapped on the phylogeny with white rectangles. mRNA loss of *UVRh2* was followed by pseudogenization in the silvaniform clade but not in the *H. melponene* clade. Protein expression presence or absence is shown to the right for males and females, presence with filled-in squares, absence with empty squares. The antibody for UVRh1 stains *Eueides isabella* UVRh, indicating antigen conservation with UVRh1. **D)** Spectral sensitivities of the UV1 and UV2 photoreceptor cells from *H. erato*, and the UV1 photoreceptor cells from *H. ismenius* (silvaniform), and *H. melpomene.* In both males and females of *H. erato,* UV2 cells have peak sensitivity at 390 nm, while only females also have the UV1 cell, whose peak is at 355 nm. *H. ismenius* (n =1) and *H. melpomene* (n=3), which both lack UVRh2 protein expression, only have a UV1 cell, which peaks at a slightly longer wavelength of 365 nm compared to *H. erato.*

The genus *Heliconius* has undergone an adaptive radiation throughout the Neotropics, driven in part by a spectacular diversity of aposematic wing patterns that also serve as sexual signals (Brown, 1981; Estrada and Gilbert, 2010; Estrada and Jiggins, 2002; Hill et al., 2013; Jiggins et al., 2001; Joron et al., 2006; Kronforst et al., 2007; Merrill et al., 2019, 2015; Smiley, 1978; Williams and Gilbert, 1981). This complex visual ecology within the genus is reflected in significant variation among the adult compound eye retinal mosaics found across species. Much of this eye diversity stems from an UV opsin (*UVRh*) duplication that occurred at the base of the genus, leading to two distinct UV-sensitive R1 and R2 photoreceptor cell types (Briscoe et al., 2010). Subsequently, lineage-specific variation in the spatial expression of particular UV-sensitive R1 and R2 cell types gave rise to at least three forms of sexual dimorphism (McCulloch et al., 2017). In the sexually dimorphic visual system of *Heliconius erato*, females express UVRh1 and UVRh2 opsins while males only express UVRh2 (Figure 1B). Intracellular recordings and behavioral experiments showed that *H. erato* females have two physiologically distinct UV photoreceptor cell types, and that these conferred true UV color vision in females only (Finkbeiner and Briscoe, 2020; McCulloch et al., 2016a). Differences in the retinal mosaics across species and sexes showcase *Heliconius* as an evolutionary model to study incipient visual system divergence (McCulloch et al., 2017).

In two other groups of *Heliconius* species, protein expression of one of the UV opsins was lost: the silvaniform clade (e.g., *H. ismenius, H. hecale*) and the *melpomene/cydno* clade do not express UVRh2 in their retinal mosaics (McCulloch et al. 2017) (Figure 1B,C). In the silvaniforms, *UVRh2* is currently undergoing pseudogenization. Independent loss-of-function mutations are accumulating in different species, suggesting this process began in parallel after the split from the *melpomene/cydno* sister clade (McCulloch et al., 2017). Meanwhile in *melpomene/cydno* species, expression of full-length *UVRh2* is low but not absent, even though protein expression is missing from the eye. Consistent with a loss of the UV2 R1 and R2 cell, our behavioral experiments indicate that both male and female *H. melpomene* are unable to discriminate between 380 nm and 390 nm light (Finkbeiner and Briscoe, 2020). Thus, it seems likely that the silvaniforms downregulated *UVRh2* expression earlier than *H. melpomene* did. Our previous work suggested that the *Heliconius* visual system (as exemplified by *H. erato*), likely compensated after neofunctionalization of *UVRh2* via spectral tuning to accommodate a new color channel (Finkbeiner and Briscoe, 2020; McCulloch et al., 2016a). As divergence went on, how did the subsequent *loss* of this same receptor in some lineages affect spectral tuning of their remaining photoreceptors? Since the loss of this UV cell type could mean the loss of color vision over a large swath of short wavelengths, we were interested in comparing the molecular, cellular, and physiological basis of this loss between these species.

To better understand the mechanisms of color vision evolution in these closely related species, we investigated opsin expression and photoreceptor cell function in *H. melpomene* and *H. ismenius.* We compared our new findings to previously published *H. erato* data, giving a unique perspective on photoreceptor cell evolution after loss of a recently gained opsin-based receptor (~4.5 Mya (Kozak et al., 2015)). We also investigated potential *cis-*regulation in the two *UVRh* loci in *H. melpomene.* Here we show that *H. ismenius* and *H. melpomene* have compensated for *UVRh2* loss via spectral tuning of other photoreceptors and identify potential transcription factors that regulate *UVRh* expression in *H. melpomene.* We also identify a novel broadband cell type in these species. Together our comparative study reveals how visual system diversity arose via both gain and loss of a UV-sensitive photoreceptor.

## Results

### UV photoreceptor spectral tuning and cis-regulation in Heliconius UVRh loci

We first asked whether UVRh2 loss might alter UV photoreceptor cell spectral sensitivity in *H. ismenius* and *H. melpomene* relative to *H. erato.* Both *H. ismenius* and *H. melpomene* lack a UVRh2-expressing cell (Figure 1C). This suggests a loss in these two species of the 390 nm cell type found in *H. erato*. Behavioral tests of *H. melpomene* confirm adult males and females are unable to discriminate 380 nm from 390 nm light (Finkbeiner and Briscoe, 2020). Intracellular recordings reported here reveal a single UV sensitive R1 and R2 photoreceptor cell type in both *H. melpomene* and *H. ismenius*, with peak sensitivity or λ_max_ = 365 nm (Figure 1D), consistent with behavioral experiments. This peak is shifted 10 nm toward longer wavelengths compared to the *H. erato* UV1 cell type with λ_max_ = 355 nm.

This loss of the UV2 cell results in a significant change in the sensory capability of *H. melpomene* (and presumably in *H. ismenius*): loss of UV color vision. To understand potential mechanisms for this loss we further investigated the molecular evolution and regulation of *UVRh1* and *UVRh2.* To test whether any sites in silvaniform/*melpomene UVRh1* are under positive selection, we performed a branch-site test of this clade against other *Heliconius* sequences, but no significant differences were detected (Table S1, Data S1). We did not test for adaptive evolution of the *UVRh2* locus in silvaniform*/melpomene* clades because *UVRh2* is a pseudogene in *H. ismenius* and *UVRh2* expression is downregulated in *H. melpomene* (McCulloch et al., 2017).

To determine if the difference in gene expression of *UVRh1* and *UVRh2* in *H. melpomene* is due to differences in *cis*-regulation, we used Assay for Transposase-Accessible Chromatin using sequencing (ATAC-seq). We targeted *H. melpomene* for sequencing due to its well-annotated genome (*Heliconius* Genome Consortium, 2012; Davey et al., 2016). We generated ATAC-seq libraries from brain and compound eye photoreceptor cells of 2 *H. melpomene* adults. Each library had on average about 35 million reads and approximately 92 percent of the reads mapped and paired properly (Table S2). We used macs2 to identify ATAC-seq read peaks denoting open chromatin and merged replicates to find consensus peaks. The *UVRh1* locus is indeed more open in general. We identified three peaks upstream of *UVRh1* in both brain and eye, and two peaks downstream one of which is only found in the eye but not brain (Figure 2A). There were no peaks near *UVRh2* in either sample, indicative of a more closed chromatin state in this region (Figure 2A). The more open chromatin state at the *UVRh1* locus is likely related to its increased expression relative to *UVRh2*. Next, we sought to understand what specifically might be regulating expression at the *UVRh1* locus.

**Figure 2.**
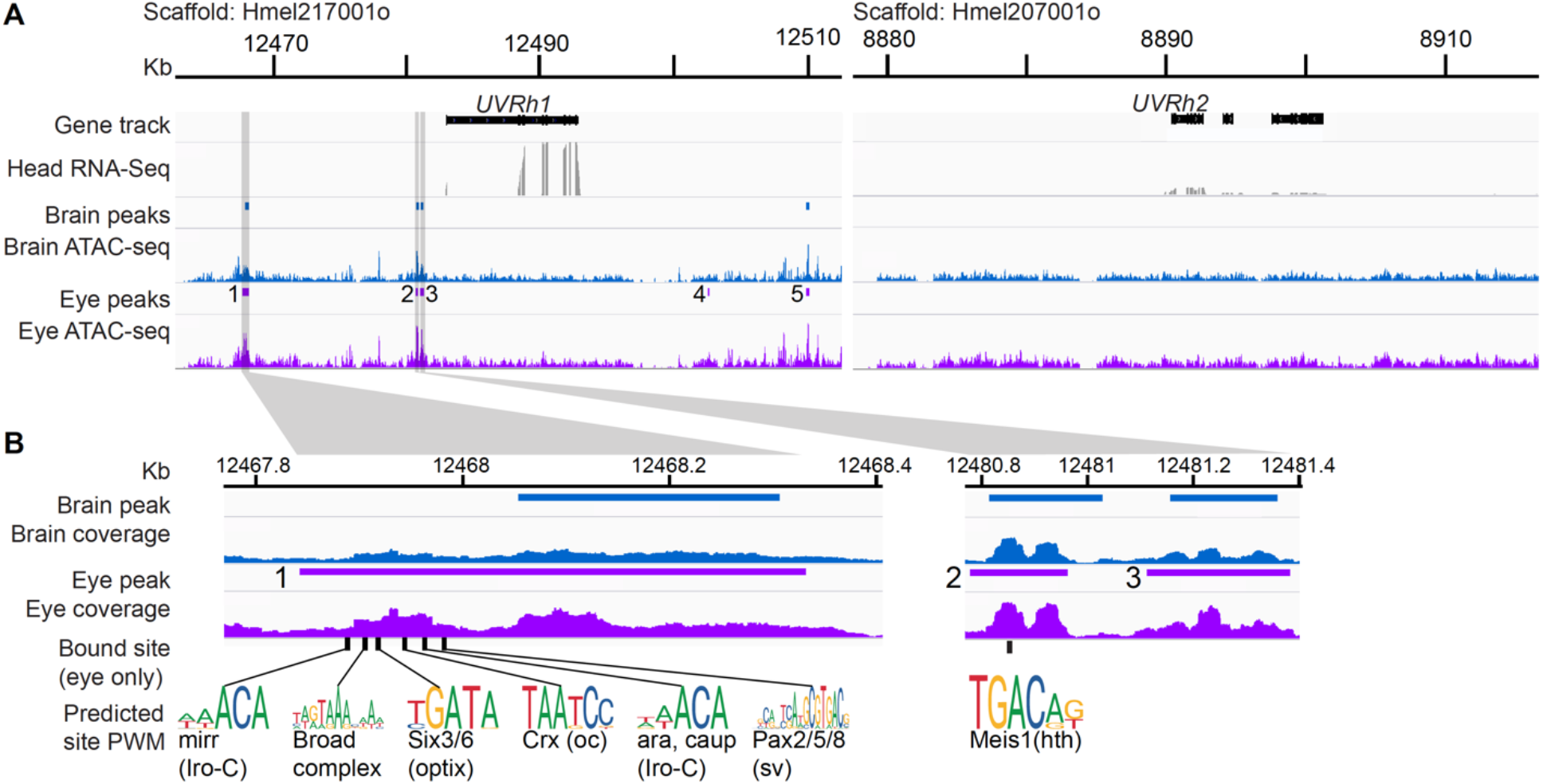
Chromatin state of *UVRh* loci in adult *H. melpomene* eye and brain. **A)** 50 kb *H. melpomene UVRh1* and *UVRh2* loci showing ATAC-seq reads and predicted peaks of open chromatin for brain (blue) and eye samples (purple), and whole-head RNA-seq reads. Five significant peaks of open chromatin are found in *UVRh1* and only four of these are significant in the brain sample. The *UVRh2* locus is much more closed and no significant peaks were detected in any sample. **B)** Zoomed in peaks showing selected transcription factor binding sites predicted to be bound by TOBIAS analysis in the *UVRh1* locus in eyes only. No brain peaks have bound transcription factors (See Table S3 for bound/unbound transcription factor binding sites in each peak). A subset of bound sites for known eye-related transcription factors are highlighted and the position weight matrices (PWMs) for each are shown (from JASPAR).

While chromatin could be open, transcription factors may not be currently bound and regulating transcription. We looked for evidence of bound transcription factors at known binding motifs using TOBIAS (Bentsen et al., 2020) in the *UVRh1* peaks in both eye and brain (Figure 2A,B, Table S3). TOBIAS uses small differences in read coverage at transcription factor binding sites to identify the site as either in the bound or unbound state (Table S3). Our data show that transcription factors are bound in the eye but not in any *UVRh1* peaks in the brain, indicating more transcriptional activity in the eye. In the eye, peaks 1, 2, and 5 have bound transcription factors. Eye peak 1 has several bound sites for transcription factors known to be involved in eye development. We identified bound sites for two members of the retinal determination gene network, Pax2/5/8 and Optix, which are classical regulators of eye development in both *Drosophila* and vertebrates (Figure 2C, Table S3) (Fu and Noll, 1997; Kumar, 2009). We also detected binding for the three TALE-class homeodomain proteins of the Iroquois complex (Araucan, Caupolican, and Mirror), which are important in eye imaginal disc development as well as adult regulators of rhodopsin expression (Mazzoni et al., 2008). Other bound sites include sites for Ocelliless/Crx, which is a paired domain homeobox protein and known regulator of rhodopsin in *Drosophila* (Tahayato et al., 2003), and the Broad-complex zinc finger, which is required for morphogenetic furrow progression and proper R8 specification in the compound eye (Brennan et al., 2001). A bound Maf subfamily site in eye peak 2 is predicted to be similar to neural-retina specific leucine zipper protein (NRL), which has been shown in mice to directly bind to the rod-opsin promoter and is required for rod differentiation, though its role in *Drosophila* eye development is unknown (Mears et al., 2001). This peak also contains a bound site for the TALE-class homeodomain protein homothorax/Meis1, involved in delimiting the eye field and in differentiation of dorsal rim area photoreceptors in *Drosophila* (Fig. 2C) (Pai et al., 1998; Wernet et al., 2003). Eye peak 5 is rich in bound sites for C2H2 zinc finger and bHLH factors. These are general transcriptional regulators and could be responsible for maintaining open chromatin at this site. Together these data suggest that *cis-*regulatory mechanisms have resulted in a more closed chromatin state at the *UVRh2* locus, blocking its expression. Although *UVRh1* chromatin is open in both the eye and brain, evidence that this locus is transcriptionally active via bound transcription factors is only found in the eye. Butterfly homologues of visual transcription factors characterized in *Drosophila*, which are expressed in the adult *H. melpomene* eye, may be involved in adult UV cell maintenance.

### Spectral tuning in LWRh- and BRh-expressing cells in H. melpomene and H. ismenius

We next investigated if the visual systems of *H. melpomene* and *H. ismenius* might have compensated for the loss of the 390 nm photoreceptor cell by tuning other spectral classes of photoreceptor. *H. erato* has a green-sensitive photoreceptor that peaks at λ_max_ = 555 nm, while this cell in *H. melpomene* and in *H. ismenius* peaks at λ_max_ = 570 nm, a shift toward longer wavelengths (Figure 3A). Purifying selection was previously found to have occurred in *Heliconius LWRh* orthologs relative to outgroup genera (Yuan et al., 2010). Similarly, we did not find positively selected sites in *melpomene/*silvaniform *LWRh* relative to the rest of *Heliconius*. In *Heliconius*, these LW photoreceptor cell types are the most common photoreceptor cells in the retina (Figure 1B) and are also the most abundant cell type found in our newly reported recordings (Figure S1, Table S4). High numbers of replicates allow for fine resolution of the spectral sensitivities in LW cells. All three species have a shoulder in the blue part of the spectrum where the cells deviate from the idealized rhodopsin curve (Figure 3A). In addition, *H. melpomene* and *H. ismenius* but not *H. erato* have a second lower peak at 400 nm in this cell type. This might be due to electrical coupling to other blue and UV photoreceptor cell inputs, or some other unknown modulation.

**Fig. 3.**
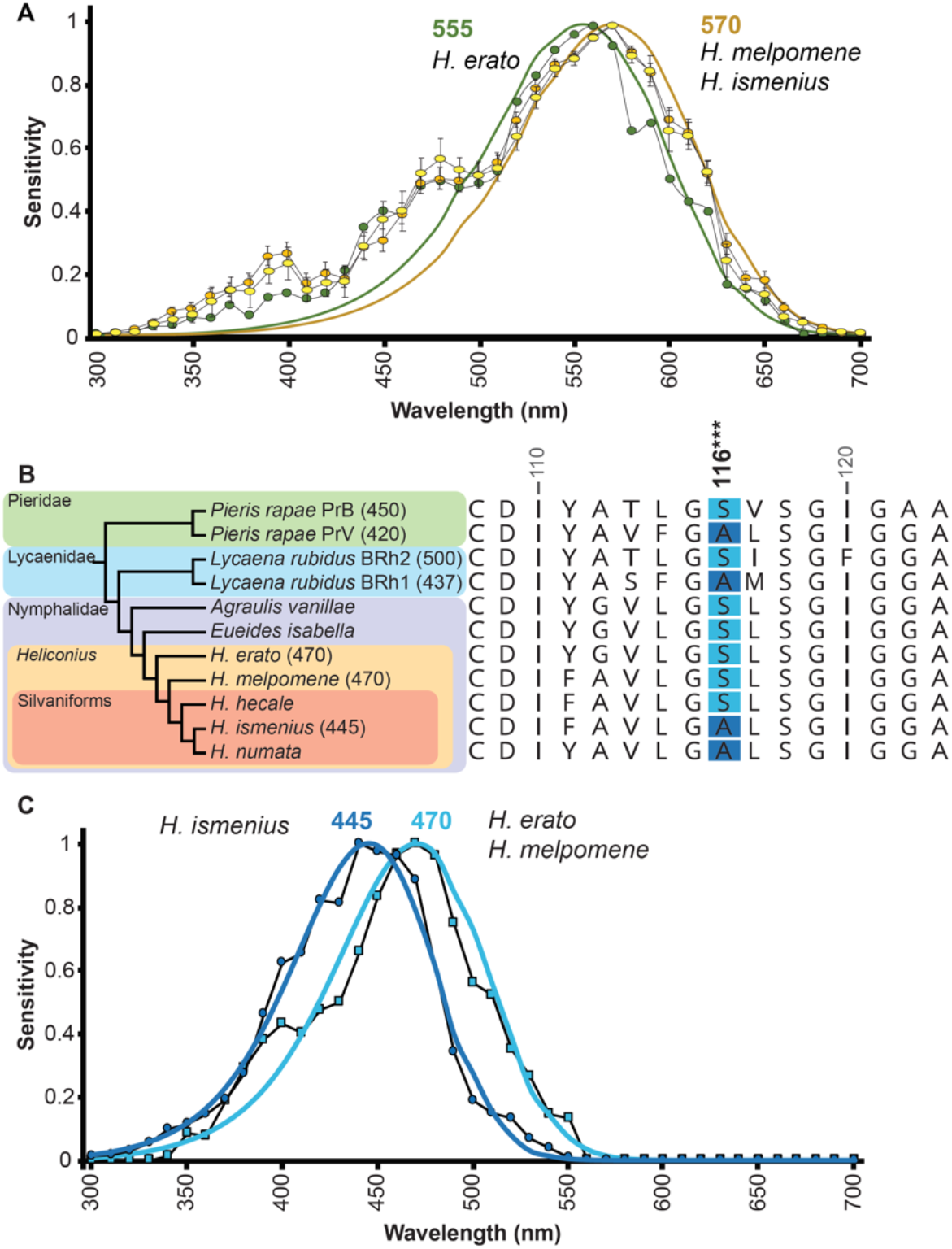
Opsin-based spectral tuning of long- and blue-wavelength photoreceptor cell sensitivities. **A)** Long-wavelength photoreceptor sensitivities of *H. erato* (n = 19) (McCulloch et al., 2016a)*, H. ismenius* (n = 11) and *H. melpomene* (n = 22) peak in the green range, at 555 nm (green circles) and 570 nm (yellow circles). **B)** Species phylogeny of butterflies (left) with known opsin peak absorbances based on intracellular recordings and retinal densitometry (Wakakuwa et al. 2010; Bernard and Remington, 1991) given in parentheses. Blue opsin alignment in the region around the spectral tuning site 116 (right). In distantly related butterflies (*Pieris, Lycaena*), a Ser116Ala substitution causes a blue-shift in blue opsin absorbance (Liénard et al., 2020; Sison-Mangus et al., 2006; Wakakuwa et al., 2010). *Heliconius* have a single blue opsin. *H. erato, H. melpomene,* and *H. hecale* have Ser at position 116 while *H. ismenius* and *H. numata* have Ala. Numbering is relative to squid rhodopsin. **C)** Averaged spectral sensitivities of *Heliconius* blue-sensitive cells. The *H. ismenius* blue-sensitive photoreceptor λ_max_ is blue-shifted to 445 nm (n = 4) relative to the *H. melpomene* (n = 7) and *H. erato* λ_max_ at 470 nm (n = 11) data not shown but see McCulloch et al. 2016.

We predicted middle-wavelength differences might exist in the absence of the UV2 R1 and R2 cell. The *H. erato* blue cell is long-wavelength shifted (λ_max_ = 470 nm) compared to most insect blue photoreceptors (~450 nm, (Briscoe and Chittka, 2001); see also van der Kooi 2013), perhaps to accommodate the presence of the 390 nm receptor. We therefore hypothesized that the blue receptor might be shifted toward shorter wavelengths in *Heliconius* species lacking the 390 nm receptor, to better discriminate colors in the blue/violet range. In *Pieris rapae*, a Ser116Ala mutation is solely responsible for a ~13 nm blue-shift in the peak absorbance of blue rhodopsin expressed in cell culture, from 450 nm to 437 nm (a Phe177Tyr substitution is responsible for a 4 nm blue-shift) (Frentiu et al., 2015; Liénard et al., 2020; Wakakuwa et al., 2010). The substitution at site 116 is present in one of the duplicated blue opsins in *P. rapae* and in the independently duplicated blue opsins of *Lycaena rubidus,* contributing to the difference between the two blue opsins’ peak absorbance within each species (Figure 3B) (Sison-Mangus et al., 2006; Wakakuwa et al., 2010). We found that *H. erato* and *H. melpomene* have serine in this position while the silvaniforms *H. ismenius* and *H. numata* have substituted Ser116Ala at this position (all *Heliconius* are fixed for Phe at site 177) (Figure 3B). This known spectral tuning site may be responsible for the long-wavelength shifted *H. erato* blue cell and suggests that *H. melpomene* also has a longer wavelength-sensitive blue cell, while *H. ismenius* may have a blue-shifted blue cell. As predicted by this spectral tuning site, intracellular recordings show that *H. melpomene* blue cells’ peak sensitivity λ_max_ = 470 nm (identical to *H. erato’s*). The *H. ismenius* blue cell peak sensitivity is shifted 25 nm toward shorter wavelengths at λ_max_ = 445 nm (Figure 3B). Other spectral tuning sites are likely also to be involved; however, the only positively selected site was the Ser116Ala as identified by Bayes Empirical Bayes inference as implemented in PAML (Table S1, Data S1) (Yang et al., 2005). Relative to *H. erato,* these results indicate *H. melpomene* and *H. ismenius* are spectrally tuning their middle and long-wavelength photoreceptors—a change coinciding with a loss of the UV2-expressing R1 and R2 cell in these species.

### Non-opsin based spectral tuning in Heliconius

Although most recordings in these species are well explained by corresponding opsin expression and known models of opsin-based absorbance spectra (Stavenga, 2010; Stavenga et al., 1993), a subset of recordings are not so easily explained. One of these is the narrow-peaked red receptor. Rare among insects (Briscoe and Chittka, 2001; van der Kooi et al., 2013), *H. erato* can discriminate red colors while another nymphalid butterfly, *Vanessa atalanta* cannot (Zaccardi et al. 2006). A red-sensitive photoreceptor cell type has been identified in *H. erato* despite no evidence for a corresponding red-absorbing rhodopsin (Bernhard et al., 1970; McCulloch et al., 2016a; See also Swihart, 1972 for recordings from red-sensitive visual interneurons). Previous work in *H. erato* found an orange filtering pigment adjacent to the rhabdom in a subset of LWRh-expressing R5-8 cells, which absorbs blue light and shifts the cells’ overall sensitivity from green to red light (Figure 1A,B) (Zaccardi et al., 2006). We wanted to know whether *H. melpomene* and *H. ismenius* also have a red-sensitive photoreceptor cell in addition to a green-sensitive photoreceptor cell. Like *H. erato,* both *H. melpomene* and *H. ismenius* do indeed have a red-sensitive cell, and in all three species the red-sensitive photoreceptor cell peaks at λ_max_ = 590 nm (Figure 4A). The red-sensitive cell thus appears to be common to all *Heliconius* species studied so far. Spectral tuning of this novel photoreceptor has proceeded through a filtering mechanism rather than through gene duplication and molecular evolution of opsin.

**Fig. 4.**
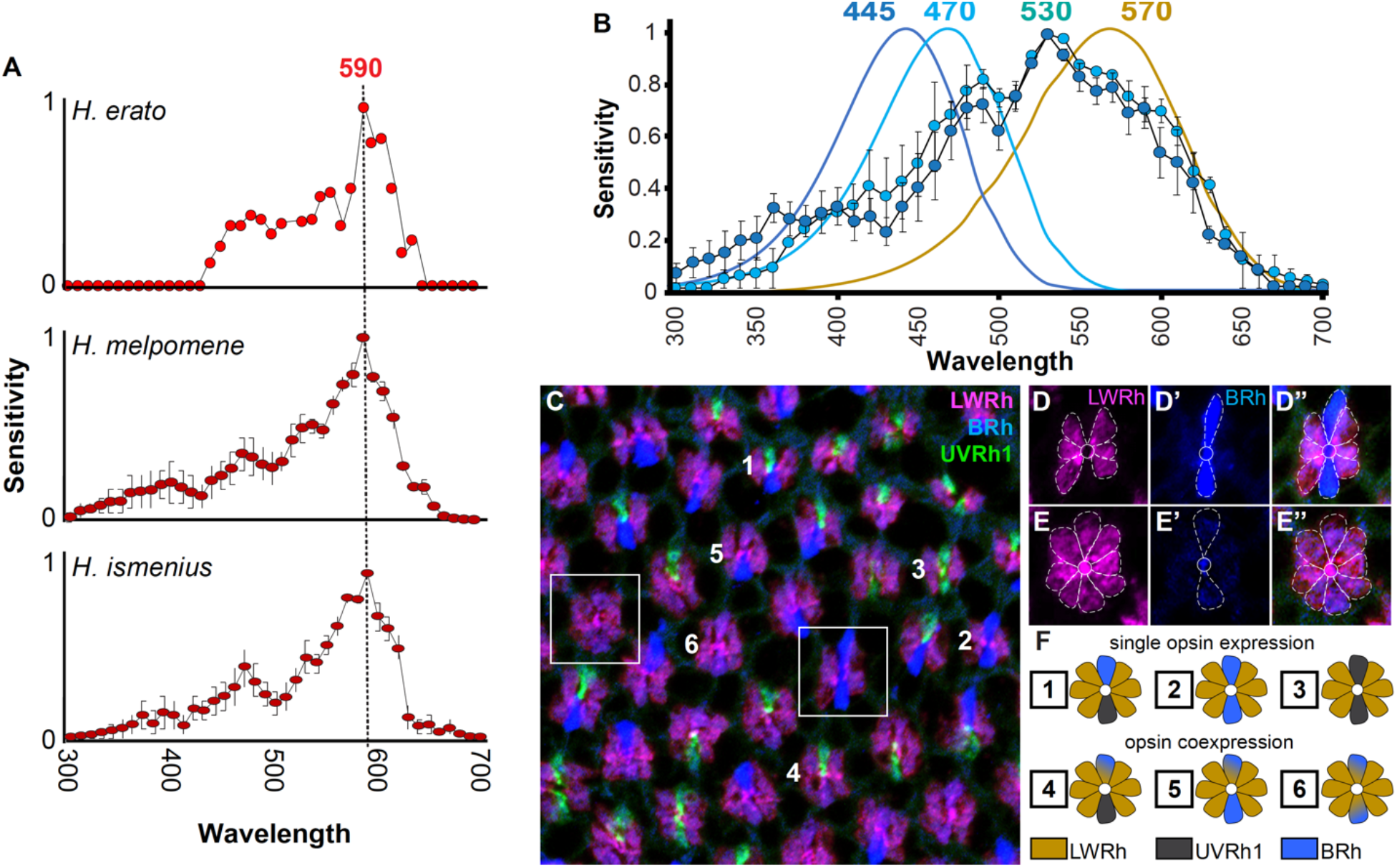
Non-opsin-amino-acid-based spectral tuning in *Heliconius* includes filtering pigments and opsin coexpression. **A)** *H. erato, H. ismenius* and *H. melpomene* all have a red-sensitive cell, which has a more narrow spectral sensitivity than rhodopsin alone and peaks at 590 nm (*H. ismenius* n = 9, *H. melpomene* n = 6). **B)** In both *H. ismenius* (n = 5) and *H. melpomene* (n = 2), we identified a cell with broad sensitivity across blue and green wavelengths, peaking at ~530 nm. The shape of the spectral sensitivity curve of this photoreceptor does not resemble any single visual rhodopsin in *Heliconius* nor does it fit with estimated opsin peaks in these species. **C-E”)** We identified LWRh in R1/R2 cells in a subset of ommatidia where BRh is also weakly expressed. Overlay of triple stain with BRh, LWRh, and UVRh1. Boxes of two example ommatidia are enlarged and cells are outlined in **D-E”**. **F)** *H. melpomene* has six ommatidial types with respect to R1/R2 expression, numbered in **C**.

Another photoreceptor found in males and females of both *H. ismenius* and *H. melpomene* is characterized by an unusually broad spectral sensitivity in the blue-green range (Figure 4B). The long wavelength shoulder roughly aligns with the 570 nm rhodopsin template, while the short wavelength (< 400 nm) region matches the blue rhodopsin template for each species. We did not find any evidence for a single opsin that might be responsible for this cell type and hypothesized that the unusually broad sensitivity could be due to opsin co-expression. However the dip in sensitivity between this photoreceptor’s peak wavelength near 530 nm and where it aligns with the blue template < 400 nm suggests filtering over these wavelengths as well.

Using an expanded combination of antibodies, we stained and carefully observed opsin expression in *H. melpomene* compound eyes. To our surprise, we identified a subset of R1 and R2 cells that weakly express BRh and also express LWRh opsins (Figure 4C-E). The presence of LWRh in R1/R2 cells represents a novel expression domain in these ommatidia. Previously we characterized the *H. melpomene* eye as a mosaic made up of three types of ommatidia, based on 3 antibodies against UVRh1, UVRh2, and BRh opsins labelling R1/R2 cells of each ommatidium (McCulloch et al., 2017). Here we identify, in addition to the previous three ommatidial types (UV1-UV1, UV1-Blue, Blue-Blue), three novel ommatidial types in *H. melpomene* (Blue/LW-Blue/LW, Blue/LW-Blue, Blue/LW-UV1) (Figure 4D-F). This brings the retinal mosaic of *H. melpomene* up to six known ommatidial types. We conclude that the broadband blue green-sensitive photoreceptor cell measured with intracellular recordings is the same cell type as that which co-expresses the LWRh and BRh opsins. We also suggest that these broadband photoreceptors contain an as-yet-unidentified short-wavelength band pass filter distinct from the red filtering pigment, perhaps bound by a new class of R1- and R2-specific ommochrome-binding protein we recently discovered (Dang and Briscoe, unpublished observation).

Since we found this BRh and LWRh co-expressing photoreceptor cell type in both *H. ismenius* and *H. melpomene*, we checked other representatives of major clades in *Heliconius* as well as outgroup genera *Eueides* and *Dryas.* In every species, we found instances of LW/B coexpression in R1/2 cells (Figure S2). This is not due to poor specificity of our antibodies, because in all instances we also identify adjacent ommatidia with photoreceptors expressing either blue or LW opsins but not both. This novel coexpression brings the total ommatidial types in *H. sara* females up to ten. Thus, we have uncovered spectral tuning as a result of a complex interaction between opsin coexpression, likely expression level differences between species, and a blue filtering pigment. Together these mechanisms result in a broadband blue green-sensitive photoreceptor cell, which also increases the number of unique ommatidial types in compound eyes across the genus.

## Discussion

This study compares photoreceptor physiology and spectral tuning in two species of *Heliconius* that have lost the UV2 (but retained the UV1) R1 and R2 receptor, relative to *H. erato* which has two UV possible photoreceptor types in R1 and R2. The full range of averaged photoreceptor spectral sensitivities for *H. ismenius* and *H. melpomene* is shown in Figure S1 and summarized in Figure 5A. For both of these species, we count a total of five spectrally distinct cell types. We found that both species have opsin and non-opsin based spectral tuning across the entire spectral range relative to *H. erato*. This indicates a shift in color vision in these species in response to ecological factors after the loss of the UV2 cell.

**Fig. 5.**
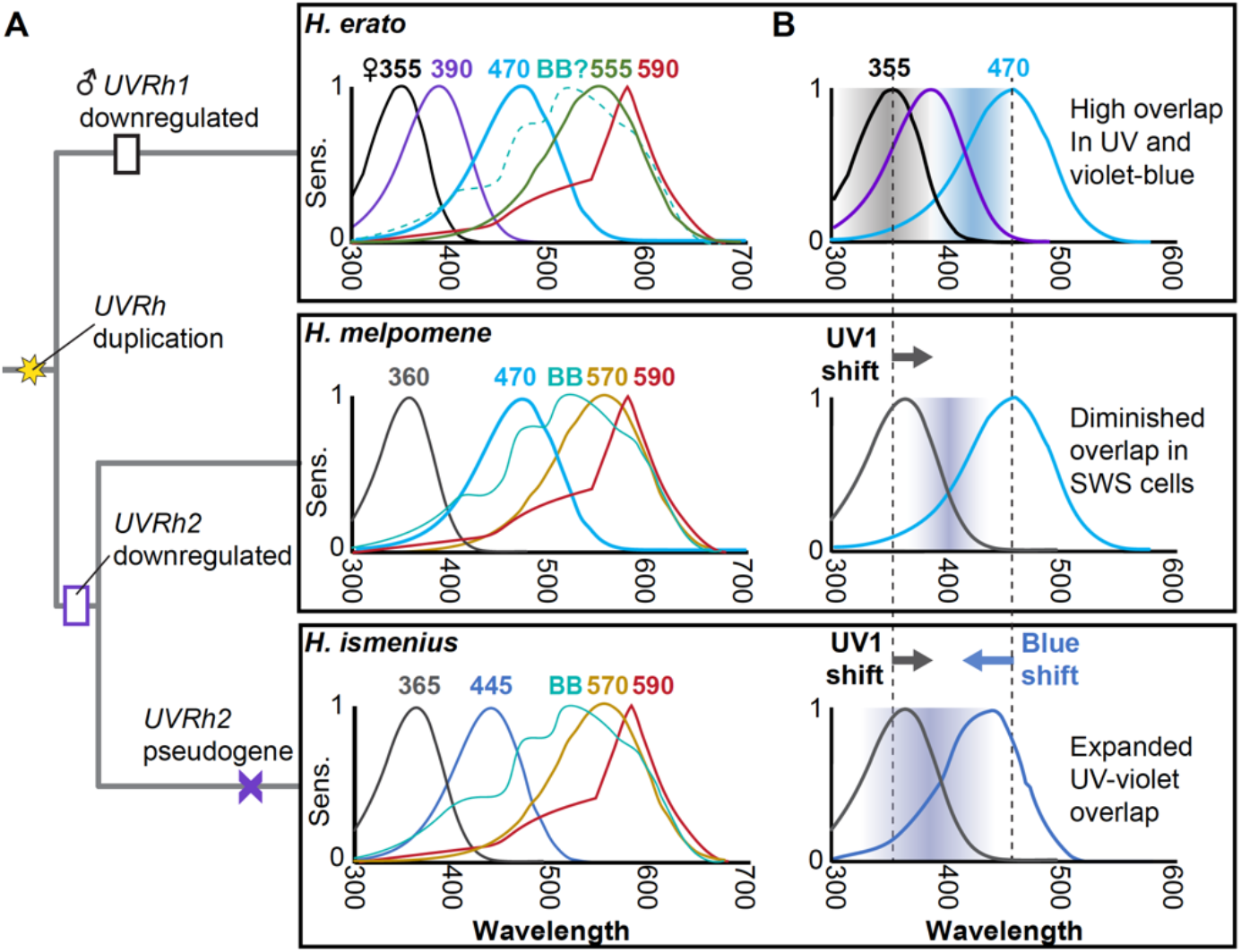
Summary of spectral sensitivities and short wavelength photoreceptor spectral tuning across three *Heliconius* species. **A)** The molecular events associated with UV opsins are mapped onto the phylogeny of the three species studied. The full spectral sensitivities of photoreceptor cells in the adult compound eye of each species are summarized. **B)** In short wavelengths, *H. erato* has significant overlap in spectral sensitivities between the two R1/R2 UV photoreceptor cell types as well as between the R1/R2 390 nm UV2 cell and the 470 nm Blue cell (shaded regions), suggesting good color discrimination in these wavelengths. In *H. melpomene* and *H. ismenius* the UV2 cell is absent, and UV1 has shifted 10 nm closer to blue wavelengths. In *H. melpomene*, the Blue cell is the same as *H. erato* resulting in diminished overlap of sensitivities in UV-to-blue wavelengths. In *H. ismenius* the Blue cell has also shifted toward the UV, and both cells now contribute to increased overlap in short wavelengths.

Our data paint a complex picture of evolutionary shifts in visual physiology in the presence or absence of the UV2 cell. We concluded from previous behavioral data that color vision in the near-UV/violet is important for female *H. erato* butterflies (Finkbeiner and Briscoe, 2020). In the absence of this violet-type receptor, shifting the peak sensitivity of the UV cell to longer wavelengths while also shifting the peak sensitivity of the blue cell to shorter wavelengths would potentially give *H. ismenius* enhanced color discrimination in this range due to greater overlap between the UV and blue photoreceptors (Figure 5B). To confirm whether spectral tuning indeed results in such a behavioral outcome, the color discrimination abilities of *H. melpomene* and *H. ismenius* in short wavelengths should be compared in the future. Establishing how the competing regimes of drift, natural, and sexual selection interplay to result in this regulatory and physiological diversity is more challenging. Tests for positive selection along branches leading to each of the silvaniform/*melpomene* opsins returned few significant sites under positive selection, suggesting drift could have had a role in this divergence, or that weak selection on multiple sites is not detected by standard methods. More behavioral and mate choice tests are necessary to better understand the ways in which the diversity of spectral receptors in *Heliconius* are used. Color signals of host plants for nectar, pollen, and oviposition may also be important and could contribute to the species- and sex-specific differences that we observe across the genus.

### Opsin molecular and genomic analysis

We wanted to identify potential spectral tuning sites for these opsins in the silvaniform/*melpomene* clade in relation to opsins in other *Heliconius* clades, so we tested each opsin for positively selected sites. None of these tests detected adaptive evolution along the silvaniform/*melpomene* branch and no spectral tuning sites were found using this approach (We did find the previously identified BRh spectral tuning site (Wakakuwa et al. 2010) via BEB inference but the likelihood ratio test between models of selection was not significant for this comparison). As has been pointed out elsewhere, adaptive variation may depend on a single amino acid substitution (Hoekstra et al., 2006), which these methods are not designed to detect (Yuan et al. 2010). Opsins are held up as examples that functional changes can be divined through their genetic code, but this study shows—using statistical tests alone—this is not always the case. With the understanding that many animal eyes are difficult to record from, we caution that using opsin mRNA expression levels and absorbance *in vitro* as a proxy for physiological sensitivity is limited. The precision and *in vivo* cellular context of intracellular recordings gives a more sensitive and biologically accurate picture of color channels that the animal is actually using.

Our *H. melpomene* ATAC-seq data corroborate our opsin expression and spectral sensitivity data for this species, where the chromatin is more open and thus actively transcribed at the *UVRh1* locus relative to the *UVRh2* locus. Peak analysis has allowed for identification of candidate transcription factors that could be maintaining UV1 cell type identity, although these are not all classical eye-related transcription factors. It has been previously shown that Pax6/eyeless binds directly to the rhodopsin promoter in *Drosophila* (Papatsenko et al., 2001; Sheng et al., 1997). A search for the consensus Pax6 *rh* binding site motif TAATYNRATTN through the *H. melpomene UVRh1* locus in the genome identifies only one motif not near any peaks in the first intron of the *UVRh1*. We also identify a Pax6 binding site in the promoter of *UVRh2*, but this is also not near a peak of open chromatin. We did not expect all eye-related developmental transcription factors to be bound to peaks in adult tissue. Many photoreceptor differentiation and determination genes may not be involved at all in adult maintenance of photoreceptor cells and adult function of developmental eye-related genes is not well-studied. Other mechanisms of transcription factor binding such as cofactor binding sites or Pax6 paired domain instead of homeodomain binding may be involved. It is therefore of note that Optix and Pax2/5/8 binding sites are bound in peak 1, suggesting combinatorial expression of particular retinal determination network paralogs could generate distinct regulatory modules in both developing and adult maintenance of *UVRh1*-expressing photoreceptor cells.

### Filtering pigments and co-expression as novel mechanisms for spectral tuning in Heliconius

Filtering of light appears to play a large role in the spectral richness of the *Heliconius* compound eye. The red-sensitive photoreceptor is invariant in all three species and displays a narrower, steeper peak than other cell types. This red sensitivity spectrum does not match a typical rhodopsin-based photoreceptor template, which predicts a much wider sensitivity range (Stavenga et al., 1993). Although the likely candidate for generating this cell type in all three species is the orange filtering-pigment identified in the *H. erato* eye (see Zaccardi et al. 2006, Fig 6C), the molecular composition of this pigment has yet to be characterized. Langer and Struwe (1972) noted the presence of an ommochrome-based pigment with peak absorbance near 560 nm in the eyes of *H. erato, H. sara,* and *H. numata*. This evidence, the location of the pigment in the eye, intracellular recordings, and behavior all support that this is the likely filtering pigment in *Heliconius*, but additional experiments are needed to confirm its identity and function. Despite distinct peak sensitivities in green cells among the three species, the composition of this unknown filtering pigment might be similar between them, because their filtered peak sensitivities are nearly identical. Furthermore, a near-infrared cell type may exist, as a visual interneuron has been reported with a peak sensitivity at 709 nm (Swihart, 1972). This could be due to signal coming from the tiny R9 cell (Figure 1A), which would only receive long wavelength light as the shorter wavelengths get filtered out by more distal receptors and filtering pigments. Alternatively, recent work has found expression of a putative retinochrome (‘unclassified’ opsin) in the crystalline cone cells and the pigment cells of the eye, which could be playing an as yet unidentified role in detection of particular wavelengths (Macias-Muñoz et al., 2019). Our intracellular recording setup currently does not presently allow us to accurately find the tiny proximal R9 cell, or still deeper interneurons, but this is an active area of further research.

Physiological evidence for a broadband cell along with immunohistochemical evidence for the co-expression of LWRh and BRh in a subset of R1 and R2 photoreceptor cells leads us to conclude that these are likely the same cell types. Our data add to growing evidence that opsin co-expression may be a common way of modulating spectral sensitivity in animal photoreceptor cells (Applebury et al., 2000; Arikawa et al., 2003; Dalton et al., 2017, 2014; Ogawa et al., 2012; Sison-Mangus et al., 2006). A broadband spectral cell type has been identified in *Papilio xuthus* via two LWRh opsins coexpressed in the R5-8 cells of a particular ommatidium, together with a 3-hydroxyretinol UV-fluorescing pigment in the distal rhabdom (Arikawa et al., 2003). Although we are unable to precisely model the *Heliconius* eye to the degree that *Papilio* has been studied, our broadband cell type spectral sensitivity is consistent with co-expression combined with a filtering pigment. Langer and Struwe (1972) identified a 450 nm maximum absorbing pigment in *Heliconius* eyes, which they speculated was a xanthommatin. We hypothesize that co-expression of BRh and LWRh leads to the observed broad, blue-shifted photoreceptor spectral peak and that filtering of blue wavelengths between 480 nm and 390 nm is occurring via this putative xanthommatin filtering pigment. Intriguingly, we have recently discovered an R1- and R2-specific ommochrome-binding protein in *H. melpomene* (Dang and Briscoe, unpublished observation). It will be interesting to see if this protein is in fact found in the subset of BRh and LW co-expressing R1/R2 cells. If this filtering mechanism is the same in *H. ismenius* and *H. melpomene*, this would also account for the fact that outside of the filtered portion of the spectra, spectral sensitivity lines up with their respective blue rhodopsin template. To confirm this, further experiments should dye inject the cell after recording, the eye should be fixed and sectioned, and then stained for these opsin proteins. We note that we also found this physiological cell type in *H. erato* but we did not report this unexplained cell type at the time. This cell type is possibly more rare in *H. erato* as blue cells are at a lower frequency overall in the retinal mosaic of this species compared to *H. ismenius* and *H. melpomene* (McCulloch et al., 2017). Finding this cell type more than once in both *H. ismenius* and *H. melpomene* was better evidence for its existence. In addition to opsin co-expression, the expression of LWRh, which is typically reserved for R3-8 cells, is found in R1/R2 cells in the tribe Heliconiini (Figure S2). This is a novel and significant finding, because it represents a regulatory switch in an otherwise highly constrained developmental program in most insects. Exceptions to this strict R1/R2 and R3-8 regulatory boundary do exist, such as in the dorsal rim area of monarch butterflies, where all R1-8 cells express UV (Sauman et al., 2005). This is the first time we observe LWRh expression in R1/R2 cells in any nymphalid.

### Conclusion

Physiological comparison of photoreceptors in the eyes of three related species shows that ongoing visual system evolution is occurring relatively quickly in *Heliconius*. We posit that after loss of the UV2 photoreceptor in *H. ismenius* and *H. melpomene,* spectral tuning of rhodopsins in these two species compensated for their loss of shorter UV color discrimination ability relative to the *H. erato* visual system. We show that a striking variety of mechanisms of spectral tuning are employed within single species in this genus, generating a diversity of spectrally distinct photoreceptors not limited by the number of opsins in the genome. Together these data showcase *Heliconius* as a remarkable example of visual system evolution following both gain and loss of a spectral channel in a colorful environment.

## Materials and Methods

### Animals

We obtained *H. ismenius telchinia* and *H. melpomene rosina* pupae from The Butterfly Farm – Costa Rica Entomological Supply. After eclosion, butterflies were housed for at least one day in a humidified chamber, and were fed a diluted honey solution daily before recording. Animals were killed after recording by rapidly severing the head and crushing the thorax.

### Opsin sequence analysis

For branch-site tests of positive selection, nucleotide alignments of *UVRh1*, *BRh1*, and *LWRh* were made from sequences published in (McCulloch et al., 2017). A maximum likelihood tree was made for each of three opsin alignments using RAxML default settings in Geneious v.9.1.5. The trees were exported as Newick files, and the branch leading to the silvaniform/*melpomene* clades was annotated as the foreground branch in the Newick file. Branch-site tests of positive selection for each tree and alignment were performed using codeML in PAML v.4.8a were compared. For the alternative model the following parameters were estimated: Ts/Tv ratio (fix_kappa = 0) and the dN/dS (fix_omega = 0), and for the null model the dN/dS ratio was set to fixed at 1 (fix_omega = 1). A likelihood ratio test was performed using the resulting likelihood scores from the alternative and null models, and significance was assessed using a χ^2^ distribution with one degree of freedom.

### H. melpomene ATAC-seq

After sacrificing, the heads of 1 male and 1 female recently eclosed *H. melpomene* were placed in a small Petri dish with Butterfly Ringer’s solution (35 mM NaCl, 36 mM KCl, 12 mM CaCl_2_, 16 mM MgCl_2_, 274 mM glucose, and 5 mM Tris-HCl, pH 7.5) to dissect and separate photoreceptor cells from brain tissue. Dissected brain and photoreceptors were placed in separate 1.7 ml microcentrifuge tubes containing 500 μl Ringer’s. The tubes were centrifuged at 500 x g for 5 minutes at 4°C. Tissue was washed with 500 μl 1x PBS buffer and centrifuged at 500 x g for 5 minutes at 4°C. We added 100 μl of cold lysis buffer (10 mM Tris-HCl, pH 7.4, 10 nM NaCl, 3 mM MgCl2, 0.1% IGEPAL CA-630 (Sigma-Aldrich, St. Louis, MO)) and ground the cells with a pestle. We transferred the ground mixture into a Nucleospin filter (Fisher Scientific, Pittsburgh, PA) and centrifuged the column at 500 x g for 10 min at 4°C. We did a transposition reaction using 25 μl 2x TD buffer, 2.5 μl Tn5 Transposase and 22.5 μl Nuclease Free water. The reaction was incubated at 37°C for 30 min and purified using a MinElute Reaction Cleanup Kit (Qiagen, Germantown, MD). The PCR reaction consisted of 30 μl transposed DNA, 2.5 μl customized Nextera PCR primer 1 and 2.5 μl primer 2, and 30 μl Phusion DNA polymerase mix (New England Biolabs). The reaction was incubated at 72°C 5 min, 98°C 30 sec, 9 cycles of [98°C 30 sec, 63°C 30 sec, 72°C 1 min], hold at 4°C. Product was run on a gel and 100 to 500 bp fragments were size selected; recovered DNA was purified using a MinElute PCR Purification Kit (Qiagen, Germantown, MD). Two replicates for each sex and tissue type (male brain, male photoreceptors, female brain, female photoreceptors) of *H. melpomene* were sequenced using Illumina paired end 43 bp reads on a Nextgen 500 sequencer. After sequencing, adapters and low-quality base pairs were trimmed using Trimmomatic v. 0.35 with the following parameters: PE [Read1.fastq] [Read2.fastq] paired_Read1.fastq.gz unmated_Read1.fastq.gz paired_Read2.fastq.gz unmated_Read2.fastq.gz ILLUMINACLIP:NexteraPE-PE.fa:2:30:8:4:true LEADING:20 TRAILING:20 SLIDINGWINDOW:4:17 MINLEN:30. Paired reads were mapped to the Hmel2.5 genome (unpublished, lepbase.org) using bwa aln in bwa v. 0.7.8. Duplicate reads were removed using Picard tools v. 1.96 (http://broadinstitute.github.io/picard) with the following parameters: MarkDuplicates INPUT=input.bam OUTPUT=output.bam METRICS_FILE=metrix.txt REMOVE_DUPLICATES=true VALIDATION_STRINGENCY=LENIENT”. Peaks were called using callpeak in macs2 v. 2.2.7.1 with default parameters. Identified peaks in any of the samples near the *UVRh1* and *UVRh2* loci were scanned using the JASPAR database for potential transcription factor binding sites, using a 95% threshold level. Peaks across tissue replicates were merged using bedtools (v. 2.25.0) intersect to find consensus peaks and bam files were merged using samtools (v. 1.9) merge. Analysis of bound and unbound motifs was done using the pipeline in TOBIAS (Bentsen et al., 2020). After merging bed and bam files, we used ATACorrect to remove Tn5 bias, ScoreBigwig to calculate footprint scores and BINDetect to determine bound positions. ATAC-seq data was visualized in IGV.

### Intracellular recording

Detailed written and video protocols are published elsewhere (McCulloch et al., 2016b, 2016a) and are described briefly here. Only one cell from one individual was used for each biological replicate. The sex of the individual was determined then it was affixed inside a small humidified plastic tube using hot wax. The tube was mounted on a stage and an 0.125 mm diameter indifferent silver electrode was inserted into the head via the mouthparts. A small hole (~10 ommatidia in diameter) was cut in the cornea using a thin razorblade and sealed with Vaseline to prevent desiccation. We used an Oriel Xenon Arc lamp (Irvine, CA, USA) as a light source. The light was passed through a condenser lens assembly (Model 60006, Newport, Irvine, CA, USA), a convex silica lens (SPX055, Newport), a neutral density (ND) filter wheel (0 to 3.5 optical density), 10 nm bandwidth spectral interference filters (40 filters, spanning 300-700 nm, Edmund Optics, Barrington, NJ, USA), a concave silica lens (Newport SPC034), a shutter with drive unit (100-2B, Uniblitz, Rochester, NY, USA), and a collimating beam probe (77644 Newport), into an attached UV-transmitting 600 μm diameter fiber optic cable (78367 Oriel), on an optical rail. Photoreceptors were recorded intracellularly with sharp borosilicate capillary microelectrodes filled with 3M KCl (~100 MΩ tip resistance). To be analyzed, the recording had to be stable throughout data collection, i.e. no change in resting potential, at least 10:1 signal to noise ratio, and large depolarizing responses (at least ~50 mV response amplitude). Responses of narrow-band spectral flashes of 50 ms were recorded, at 0.5 s time intervals and covering the spectrum from 300 to 700 nm in steps of 10 nm. Intensity response curves were recorded from 3.5 to 0 optical density before and after an experiment to confirm no change in the recording throughout.

Spectral sensitivities of cells were derived as follows. The responses to white light at each ND filter step were used to create a response–log intensity (*V*log*I*) curve. The *V*log*I* data were used to estimate parameters for the Naka–Rushton equation: *V/V*_*max*_=*I*^*n*^/(*I*^*n*^+*K*^*n*^), where *V* is the amplitude of a given response; *V*_*max*_ is the maximum response amplitude; I is the intensity of the stimulus for the given response, *V*; *K* is the intensity of the stimulus that elicits half of *Vmax*; and *n* is the exponential slope of the function (Aylward, 1989; McCulloch et al., 2016a, 2016b; Naka and Rushton, 1966). Correction factors were calculated to approximate constant photon flux over all filters from 300 to 700 nm to account for differences in total photon flux for each interference filter. Corrected intensities were divided by the maximum response intensity for each cell to calculate relative spectral sensitivity. Photoreceptors were classified by peak sensitivity and shape of the spectral sensitivity curve and replicates were averaged and standard error was calculated for the sensitivity at each wavelegnth. To estimate peak sensitivities, we used least-squares regression to fit rhodopsin templates to our data (Stavenga, 2010). All cells and sensitivities are found in Table S4 and Figure S1.

### Cryosectioning and immunohistochemistry

Detailed methods and antibody generation are described in (McCulloch et al., 2017, 2016a). Briefly, freshly severed butterfly eyes were immediately fixed in 4% paraformaldehyde (Electron Microscopy Sciences, Hatfield, PA, USA) in 1x phosphate-buffered saline (PBS) for 30 min at room temperature. Eyes were then step-wise sucrose-protected up to 30% in PBS. Each eye was placed in Tissue Tek O.C.T. compound (VWR, Radnor, PA, USA), frozen at −20°C, and sectioned at 14 μm thickness on a Microm HM 500 OM microtome cryostat (Fisher Scientific, Pittsburgh, PA, USA). Slides were dried overnight at room temperature. Dry slides were placed in 100% ice-cold acetone for 5 min, then washed 3×10 min in PBS. Slides were then placed in 0.5% sodium dodecyl sulfate in PBS for 5 min. Each slide was blocked for 1 h at room temperature using 8% (v/v) normal donkey serum and normal goat serum, and 0.3% Triton X-100 in PBS. Slides were incubated with 1:15 guinea pig anti-UVRh1, 1:15 rat anti-BRh and 1:15 rabbit anti-LWRh antibody in blocking solution overnight at 4°C. Slides were washed 3×10 min in PBS and then incubated with 1:1000 goat anti-rat Alexafluor 488 and 1:500 donkey anti-rabbit Alexafluor 555, and 1:250 goat anti-guinea pig Alexafluor 633 (Life Technologies) in blocking solution for 2 h at room temperature. Slides were washed again 3×10 min in PBS. Images were taken using a Zeiss LSM 700 confocal microscope under a 20x objective, in the UC Irvine Optical Core Facility. Stains were pseudocolored, and contrast and brightness were adjusted for clarity using Adobe Photoshop CS4 and Fiji (Schindelin et al., 2012).

## Supporting information

Supplementary_FiguresTables

## Acknowledgements

We would like to thank The Butterfly Farm – Costa Rica Entomological Supply for help with rearing butterflies and Andrew Gehrke, Primoz Pirih, and Daniel Osorio for helpful discussions. This work was funded by a Hewitt Postdoctoral Fellowship to A.M.M. and NSF grants IOS-1257627 and IOS-1656260 to A.D.B.

