## Supplementary_FiguresTables for "Multiple mechanisms of photoreceptor spectral tuning following loss of UV color vision in *Heliconius* butterflies"

**Supplementary Data**

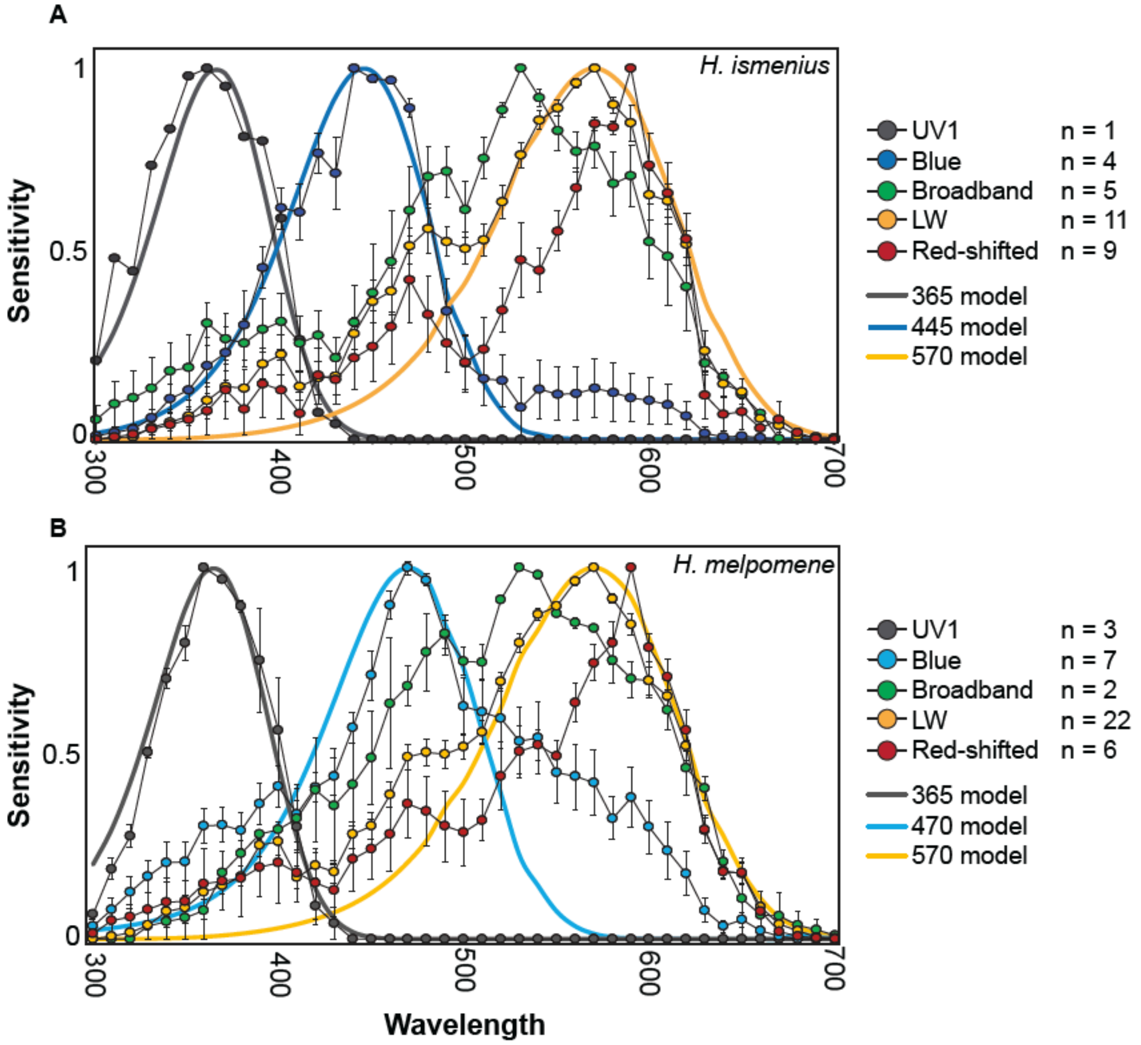


**Figure S1. Averaged spectral sensitivities in *H. ismenius* and *H. melpomene.*** All recordings used for **A)** *H. ismenius* and **B)** *H. melpomene* were averaged and presented here with standard error bars. Rhodopsin absorbance spectra are also shown for cells that appear to express a single kind of opsin. For each photoreceptor cell type, the number (n) of individual cells recorded from are listed.

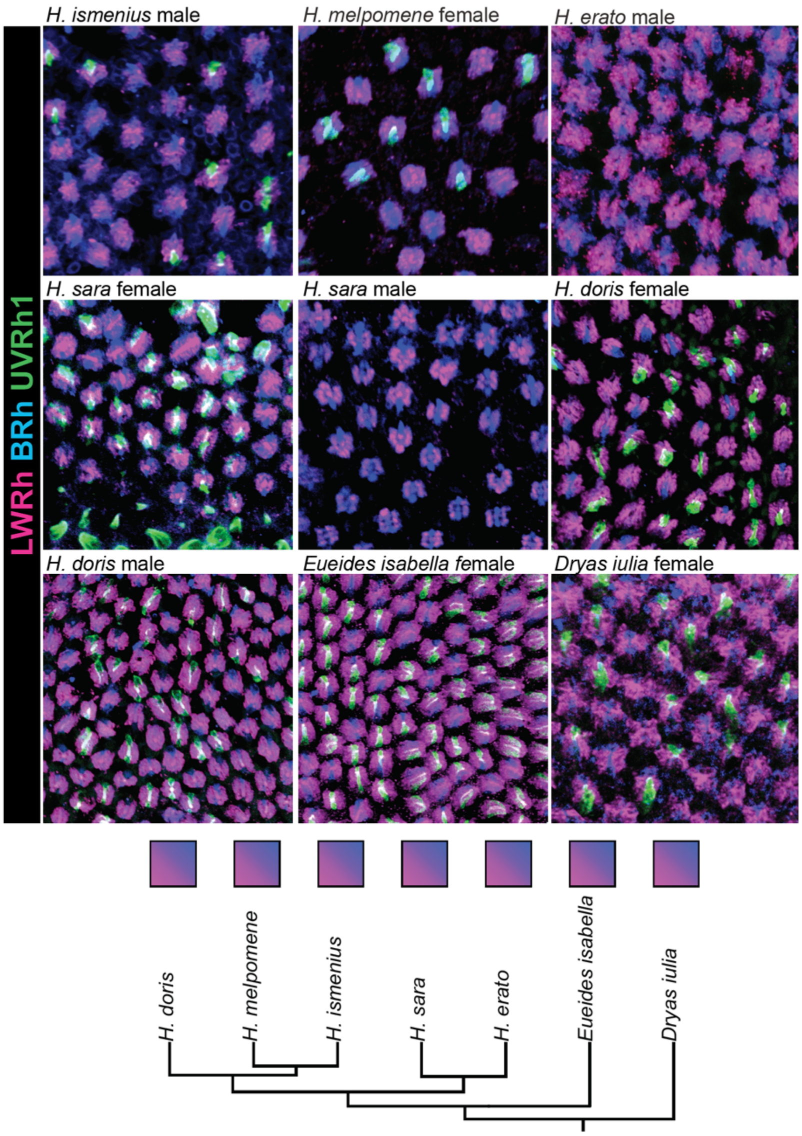


**Figure S2. Antibody stains for LWRh and BRh show opsin coexpression in *Heliconius* species and outgroups.** Triple labelling of photoreceptor cells in the adult compound eyes of *H. ismenius, H. melpomene, H. erato, H. sara, H. doris, Eueides isabella* and *Dryas iulia* using anti-LWRh (pink), anti-BRh (blue) and anti-UVRh1 (green) antibodies reveals coexpression of LWRh and BRh in every major clade within *Heliconius* as well as in the outgroup genera *Eueides* and *Dryas.*

**Table S1. PAML likelihoods for opsin branch-site tests of selection.**

| Condition | ln Likelihood | LRT | Chi-squared test p value |
|---|---|---|---|
| UVRh1_fixed | -6771.42 | 0.0600 | 0.8 |
| UVRh1_alt | -6771.39 | | |
| BRh_fixed | -4938.22 | 0.6700 | 0.41 |
| BRh_alt | -4937.88 | | |
| LWRh_fixed | -3549.31 | 0.0002 | 0.99 |
| LWRh_alt | -3549.31 | | |

**Table S2. ATAC-seq samples and statistics.**

| Sample | Specimen | Sex | Tissue | Reads | Paired | % paired | Reads Aligned | % mapped | % properly paired | Total Peaks |
|---|---|---|---|---|---|---|---|---|---|---|
| HMP516b | HMP516 | F | brain | 31518059 | 30372185 | 96.36 | 36740592 | 100 | 91.68 | 13275 |
| HMP516e | HMP516 | F | photoreceptors | 36448289 | 35180835 | 95.52 | 42110559 | 100 | 87.79 | 9009 |
| HMP517b | HMP517 | M | brain | 36025856 | 34750866 | 96.46 | 41873107 | 100 | 91.76 | 22890 |
| HMP517e | HMP517 | M | photoreceptors | 55863519 | 53884930 | 96.46 | 65897894 | 100 | 90.39 | 6304 |
| HMP540b | HMP540 | F | brain | 41305222 | 39817558 | 96.4 | 45787536 | 100 | 91.53 | 4101 |
| HMP540e | HMP540 | F | photoreceptors | 30515084 | 29270537 | 95.92 | 36601722 | 100 | 92.57 | 4781 |
| HMP541b | HMP541 | M | brain | 16349872 | 15661333 | 95.79 | 20077349 | 100 | 91.39 | 4457 |
| HMP541e | HMP541 | M | photoreceptors | 30691619 | 29773599 | 97.01 | 37971102 | 100 | 93.02 | 5455 |

**Table S3. TOBIAS footprinting scores for eye and brain *UVRh1* and *UVRh2* loci.**

(separate table)

**Table S4. Species, sex, and number of recorded cells used in this study.**

| Species | Sex | Cell Peak | | | | | |
|---|---|---|---|---|---|---|---|
| | | 365 | 445 | 470 | 530 | 570 | 590 |
| *H. ismenius* | female | 1 | 1 | 0 | 3 | 6 | 5 |
| *H. ismenius* | male | 0 | 3 | 0 | 2 | 5 | 4 |
| *H. melpomene* | female | 0 | 0 | 1 | 1 | 8 | 4 |
| *H. melpomene* | male | 3 | 0 | 6 | 2 | 14 | 2 |

**Supplementary Data** PAML alignments and trees

(separate files)
